# From overconfidence to task dropout in a spatial navigation test: evidence for the Dunning-Kruger effect across 46 countries

**DOI:** 10.1101/2023.09.18.558324

**Authors:** S. Walkowiak, A. Coutrot, J. M. Wiener, M. Hornberger, E. Manley, H. J. Spiers

## Abstract

Overconfidence and the Dunning-Kruger effect have been reported in many cognitive domains. However, there is little examination in the field of spatial navigation. Here, we examined overconfidence in navigation ability in 376,836 participants from 46 countries. We tested navigation using our virtual wayfinding task in the app-based video game Sea Hero Quest, examined self-ratings of navigation ability and how many game levels participants had played before they dropped out. The main goal of this analysis was to investigate how performance of overconfident participants influenced the dropout rate from our experimental task embedded in a video game. First, we measured and modelled overestimation at baseline game levels. Age was found to be the strongest predictor of overestimation across the entire sample. Higher age was associated with increased overconfidence in all 46 countries and 11 distinct cultural clusters. Female participants were more likely to overestimate their performance across most of the life course (19-59 years old), however older men (60-70 years old) displayed highest overconfidence amongst all age-gender groups. Overconfidence also varied widely across countries. Secondly, we estimated performance on follow-up game levels for those who were previously identified as overconfident and we found that those who were more likely to display the Dunning-Kruger effect (*i.e.*, poor performance while being overconfident) were predominantly female and older participants. Finally, survival analysis methods with time-dependent covariates revealed that poor wayfinding performance, while being overconfident, was one of the strongest predictors of task dropout. This Dunning-Kruger bias on participants dropout existed universally across all countries in our data.

## Introduction

### Defining overconfidence

Psychological research often relies on self-assessed measurements of one’s abilities. The self-reported beliefs about participants’ skills are commonly compared with either their actual behavioural performance on a variety of cognitive, motor, or psychometric tests (Lesch & Hancock, 2004; Garg et al., 2023) or with relation to their judgments about the abilities of their peers (Svensson, 1981; Alicke & Govorlin, 2005). The discrepancy between the subjective self-assessment and the objectively recorded test performance can be quantified and explained as the level of overconfidence (or underconfidence) a person has in their own task-specific abilities. Based on their Bayesian belief updating model of overconfidence, Moore & Healy (2008) pointed out that overconfidence is a multifaceted construct with three separate components which may have different impacts on actual behaviour and performance. Firstly, overconfidence may simply result from an *overestimation* of one’s skills when compared to objective performance measurements. Secondly, its source can be what Moore & Healy call an *overplacement* (also referred to as the *Above Average Effect* by Alicke & Govorlin, 2005; or the *optimism bias* by White, Eiser & Harris, 2004) – a belief that a person is more skilled or competent than their peers. Finally, overconfidence may arise due to *overprecision* in the accuracy of individual’s beliefs which manifests itself by the inability of correctly estimating the 90% confidence intervals around a particular measurement (Moore, Carter, & Yang, 2015) and it’s therefore typically defined as the excessive faith that one knows the truth. In this study, we focus on investigating overestimation, and we use terms overconfidence and overestimation interchangeably.

### From overconfidence to Dunning-Kruger effect and task dropout

Kruger & Dunning (1999) reported that the most overconfident individuals often perform poorly and largely overestimate their abilities compared to those who tend to self-assess their skills more cautiously. On the other hand, high performers display a slight underestimation of their abilities which is likely caused by their incorrect assumption about skills of others (*false consensus hypothesis*). A discussion on the exact explanation of this phenomenon, called the Dunning-Kruger effect (2011), is still ongoing, but according to Kruger & Dunning themselves, the overestimation of abilities by low performers may be caused by the failure to recognise their incompetence due to the lack of metacognitive ability. This approach has led to those who fall for this cognitive bias being labelled as *“unskilled and unaware”*. In contrast, Krajc & Ortmann (2008) proposed that the Dunning-Kruger effect is more a problem of inference or *“signal extraction”* rather than the result of biased judgments. They argued that those who perform poorly often self-assess themselves as skilled because they simply have no access to information or feedback about their own versus the others’ performance. However, this explanation was criticised by Schlösser et al. (2013) who did not find empirical evidence to support Krajc & Ortmann’s hypothesis.

The Dunning-Kruger effect received further criticism for the methodology used in typical studies reporting its existence. It is commonly estimated by either calculating average overestimation by performance quartiles or by regressing overestimation on performance. According to Feld, Sauermann, & de Grip (2017) both methods, but especially the linear regression approach, lead to biased estimates. This is due to the usually unaccounted measurement error (or *“bad luck”*) which often decreases performance and causes a substantial overestimation of the Dunning-Kruger effect in the ordinary least squares framework. Instead of that approach they propose an alternative two-stage approach correcting for this measurement bias by taking a second performance measurement as an instrumental variable. As a result of this methodological adjustment, the Dunning-Kruger effect was still present but the relationship between the performance and overestimation became significantly weaker than in the linear regression. Recently, Gignac & Zajenkowski (2020) extended their criticism even further and referred to the Dunning-Kruger effects identified in studies which implemented commonly used methods as *“(mostly) statistical artefacts”*. Using data on self-assessed and objectively measured intelligence, they tested the Dunning-Kruger hypothesis for heteroskedasticity using the Glejser test and found that the degree to which individuals overestimated their intelligence was equal regardless of the level of objective IQ. This finding contrasts with the original Dunning-Kruger hypothesis which assumes that low performers lack metacognitive ability to identify their shortcomings, and hence, if it was correct, the overestimation would be larger for these individuals rather than equal. Gignac & Zajenkowski (2020) admitted however that the hypothesis and methodology used in typical Dunning-Kruger studies may still be valid for some skills, but similarly to Feld, Sauermann, & de Grip’s (2017) conclusion, they argued that the true magnitudes of effects are likely to be smaller than commonly reported.

Despite the criticism of the applied methodology and the ongoing debate on validity of the Dunning-Kruger hypothesis, the effect has been extensively reported in studies across different fields, e.g. in academic performance and grade prediction (Magnus & Peresetsky, 2018), driving abilities and driving safety (Martinussen, Møller, & Prato, 2017), maths, physics and chemistry knowledge (Bell & Volckmann, 2011; Lindsey & Nagel, 2015; Pazicni & Bauer, 2014), self versus others assessment of intelligence (Neubauer & Hofer, 2021), second language learning (Saito et al., 2020; Stankov, Lee, & Paek, 2009), software engineering skills (Jørgensen, Bergersen, & Liestøl, 2021), personal health awareness (Miller et al., 2019), healthcare quality (Kovacs, Lagarde, & Cairns, 2020; Taylor et al., 2020), marketing and lobbying (Lyons, McKay, & Reifler, 2020), investment decision making (Lewis, 2018), sports and sports coaching (Sullivan, Ragogna, & Dithurbide, 2019), and fake news receptivity (Pennycook & Rand, 2020; Lyons et al., 2021).

Some authors have highlighted the importance of social, health and public safety implications resulting from the Dunning-Kruger effect. For example, Martinussen, Møller, & Prato (2017) warned of gross inconsistencies between self-assessed hazard prediction skills and the actual driving performance of young male drivers. On the other hand, Miller et al. (2019) pointed out that the Dunning-Kruger effect may be the main reason behind the misjudgement of health-relevant behaviours by unhealthy people and can potentially lead to their reluctance in choosing health improving interventions. Similarly, Benegal (2018) hypothesised that the excessive overconfidence and the resulting Dunning Kruger effect are the possible sources of discounting expert medical knowledge on topics such as vaccinations. Finally, according to Clark & Saxberg (2018) an overly optimistic self-assessment often leads to overconfidence and poor mental workload which may result in task disengagement, sub-optimal performance and, eventually, task failure.

### Individual differences and cultural correlates of overconfidence and the Dunning-Kruger effect

A predisposition to overconfidence and the Dunning-Kruger effect might be dependent on individual, personality, and cultural factors. Despite a popular opinion that young people are more likely to overestimate their skills and competencies, Hansson et al. (2008) found that older adults displayed higher levels of overconfidence and this age-related increase was partly mediated by general cognitive abilities. On the other hand, Prims & Moore (2017) did not report any evidence of correlation between age and overestimation, but they found a significant age-dependent increase of precision in judgement supporting the claim that older individuals may be more resistant to change their beliefs and may even become more ideologically extreme. Despite these mixed results, it is however possible that a correlation between age and overconfidence is present for some skills but not for others. For example, Taillade et al. (2013, 2016) and more recently Walkowiak et al. (2023) showed that older participants were more likely to overestimate their spatial abilities but performed worse than younger adults on wayfinding tasks. Similarly, Ulrich, Grill, & Flanagin (2019) argued that older adults have a higher propensity to overestimate their navigational ability. In the well-known study on driving performance and cell phone use, Lesch & Hancock (2004) demonstrated that the driving performance of highly confident older female drivers while using a phone was affected to a greater extent than performance of males or young females. Although the research on gender differences in task-specific confidence points towards the advantage of males, these differences in overconfidence are less obvious. Some authors argue that overestimation of abilities is universal across both men and women (Lundeberg, Fox, & Punćochaŕ, 1994), but men may display stronger tendencies for being overconfident attributing this to lower risk aversion (Soll & Klayman, 2004) and higher level of competitiveness (Reuben et al., 2012). Females were also found to report lower spatial confidence despite performing equally well as males on the reorientation task (Picucci, Caffò, & Bosco, 2011).

Cultural determinants of overconfidence, and specifically the overestimation and Dunning-Kruger effect, have also attracted rather limited scientific attention. Some of the rare examples of such investigations are two studies by Stankov, Lee & Paek (2009) and Stankov & Lee (2014). In the former, they reported that males and African American or Hispanic individuals displayed higher levels of overconfidence than females and European Americans on three cognitive tests of English language curriculum. In the latter, they showed evidence for significant cross-cultural differences in performance on a cognitive test, along with relatively small differences in confidence across 33 countries. In consequence, they reported differences in overconfidence between 9 world regions, with East Asian nations showing better calibration (*i.e.*, more accurate self-estimates when compared to the actual performance) than countries from other regions. More recently, Moore, Dev & Goncharova (2018) compared the levels of all three previously mentioned dimensions of overconfidence (*i.e.*, overestimation, overplacement, and overprecision) between the individualistic (United States and United Kingdom) and collectivistic (India and China) countries. They found that Indian participants exhibited higher overconfidence, however only with regards to overestimation and not in terms of overplacement or overprecision. In the discussion, the authors admitted that cross-cultural differences in overconfidence may be more complex in nature and are likely to be task dependent. Finally, in our earlier work (Walkowiak et al., 2023), we found that certain countries which shared cultural heritage and value systems displayed similar accuracies between their self-assessed navigation ability and the objectively measured wayfinding performance. For example, individuals from Germanic countries (Germany, Austria, and Switzerland) reported higher overestimation than representatives of the Nordic, Confucian Asia, and the Far East cultural clusters. We argued that the level of overconfidence may be partially mediated by gender stereotypes measured with Hofstede’s masculinity dimension (Hofstede, 2001; Hofstede, Hofstede, & Minkov, 2010) - greater overconfidence was associated with higher masculinity at the national level.

However, despite many studies on overconfidence across numerous research domains, none of the presented studies attempted to measure the impact of the Dunning-Kruger effect on performance in wayfinding tasks, the completion rate of multiple related tasks and the resulting retention of participants across the tests. Furthermore, most cross-cultural investigations of the Dunning-Kruger effect have been implemented on relatively small samples representing only several selected countries.

### The present study

In the current study, we seek to answer three main research questions using wayfinding data collected with the Sea Hero Quest – a mobile game application played by over 4 million users globally. While we previously reported individual and cultural differences of self-estimates, quantified and explained the gap between the self-reported navigation abilities and wayfinding performance across 46 countries and 11 cultural clusters (Walkowiak et al., 2023), we now explore cross-cultural differences and correlates of wayfinding *overconfidence* such as age, gender, and education. Secondly, we identify and describe characteristics of participants who are prone to the Dunning-Kruger effect with particular focus on those who performed poorly while displaying excessive overconfidence across multiple wayfinding tests. Finally, we investigate how overestimation of abilities and Dunning-Kruger effect impacted the risk of dropping out on subsequent spatial navigation tasks and whether this effect was present across different countries.

## Methods

### Dataset

Statistical analyses reported in this manuscript were based on samples obtained from the full behavioural and demographic dataset of 4.1 million participants who between April 2016 and April 2019 downloaded and played a mobile game application called Sea Hero Quest. The game, which comprises forty-five wayfinding, fifteen path integration, five radial maze and further fifteen strictly game engagement levels, was designed to measure typical behaviours of healthy individuals on navigation and spatial tasks across the population. During the wayfinding levels, which were of our main interest in this study, the participants had to navigate a virtual boat from the origin to the destination by following the sequence of numbered buoys which functioned as checkpoints along the route. Before each wayfinding level, a map with the layout and locations of both origin and destination points as well as positions of buoys was displayed to enable the players to plan and memorise the route. As some wayfinding levels introduced foggy/rainy conditions, parts of their maps were obscured to make it more difficult for participants to locate themselves correctly and navigate towards the intermediary buoys and final destination point. Time spent on viewing the maps was recorded and used as one of the variables in the analysis. Apart from the in-game movement and trajectory data, the participants voluntarily provided their socio-demographic information, e.g. age, gender, attained education level, country, self-estimates of their navigation skills, average daily amount of commute time, typical duration of sleep a day, and whether they were raised in rural, city or mixed areas, and which hand they used for writing. The design and experimental setup of the game with description of specific tasks has been detailed in Coutrot et al. (2018), Spiers, Coutrot & Hornberger (2023) and Walkowiak et al. (2023), whereas the evidence for ecological validity has been provided by Coutrot et al. (2019).

### Ethics and informed consent

Informed consent was obtained from all participants who played the Sea Hero Quest mobile application game. The starting screen of the game displayed debriefing information including goals of the study, what data were collected and how they were used for research and analysis. Participants could also access the full explanation of the study and the Sea Hero Quest game during the gameplay through the ‘journal’ icon. They could also withdraw from playing any part of the game or sharing their data at any point during the study.

This study was conducted as part of a larger research project which has been approved by the University College London Ethics Research Committee under the project number: CPB/2013/015. All experiments and methods including data collection, processing and analysis were carried out in accordance with relevant guidelines and regulations.

### Data cleaning procedures and sampling

The full Sea Hero Quest dataset included 4.1 million users worldwide and 50 million gameplays at forty-five wayfinding levels. Our final sample was restricted to only those players who provided self-estimates of their navigation skills, answered socio-demographic questions such as age, gender, education level, average daily commute time, typical amount of sleep and the type of home environment they grew up in, represented countries which were included in the cultural clusters identified in the *Definition of cultural clusters* subsection of Methods, and with at least 500 valid users at Level 12 (for baseline measurement of overconfidence) or at Level 38 (for estimation of dropout rate as a result of the Dunning-Kruger effect). These selection criteria were implemented to ensure a large enough sample for each country at the last task to allow statistically meaningful between-countries comparisons. Additionally, all participants had to be between 19 and 70 years old to control for fake age demographics as recommended earlier by Coutrot et al. (2018). We have also removed users who slept less than 5 and more than 12 hours a day on average. Furthermore, wayfinding performance metrics described in the *Measuring wayfinding performance* subsection of Methods were calculated using the player’s first attempts at each game level to limit the effect of training through the level repetition. We have also removed the records of participants who attempted the wayfinding follow-up levels in non-sequential order, e.g. completed level 21 before playing level 13. This was done to ensure that each participant followed the same sequence of experiments with incrementally increasing difficulty and that their performance on earlier follow-up levels was not affected by the training effect through their prior exposure to more difficult levels (or their performance on more difficult levels wasn’t compromised by their incompletion of earlier, easier levels). Finally, based on the wayfinding performance metric calculated for baseline levels (*i.e.*, defined in the *Measuring wayfinding performance* subsection of Methods), we have excluded players who performed two standard deviations below and above the sample mean on these tasks.

Following these data cleaning and processing activities, the final sample of participants who completed all baseline levels (up to and including level 12) consisted of 376,836 individuals representing 46 countries with at least 500 valid players: 172,368 females (45.74% of the overall sample; *MeanAge_Females_* = 37.86 ± 14.71) and 204,468 males (54.26%; *MeanAge_Males_* = 37.47 ± 13.48). Descriptive statistics for this sample with a breakdown by gender, country and cultural cluster are presented in Tables S1 and S2 of the Supporting Information.

Hypotheses related to the general dropout rate amongst both overconfident and not overconfident individuals were investigated using a sample of participants who were included in the sample for baseline levels and completed at least one of the wayfinding follow-up levels. Overall, 295,474 such individuals (137,442 females and 158,032 males) from 16 countries completed at least one of the sixteen follow-up levels (*MeanAge_Females_* = 38.50 ± 14.81; *MeanAge_Males_* = 38.17 ± 13.63). Descriptives of the sample cross-tabulated by gender and country are available in Table S3 of the Supporting Information.

Finally, to investigate the effects of overconfidence and poor performance (*i.e.* the Dunning-Kruger effect) on game dropout, we have selected only those participants who completed at least one of the wayfinding follow-up levels and were previously identified as overconfident at baseline levels. This sample included 86,692 participants (47,123 females and 39,569 males; *MeanAge_Females_* = 44.73 ± 14.48; *MeanAge_Males_* = 46.45 ± 13.46) from 16 countries. Table S4 of the Supporting Information displays descriptives by country and gender.

### Baseline and follow-up game levels

The Sea Hero Quest game consisted of forty-five wayfinding levels of different spatial layouts, varying conditions (e.g. foggy, clear or rainy weather, presence of sea rocks, varying height of waves etc.) and incrementally increasing difficulty (Spiers, Coutrot, & Hornberger, 2023; Yesiltepe et al., 2023). Across all levels, the players were expected to navigate a boat from the origin to the destination by following a sequence of numbered checkpoints (buoys). The locations of checkpoints and the geometric layouts of specific game levels were shown before each level as a map which the participants could study for as long as they wanted. The map view duration for each level was recorded and used later as one of the time-dependent covariates in the analysis. Due to the simplicity of levels 1 and 2, they were only used to correct the wayfinding performance metrics (see the *Measuring wayfinding performance* subsection below for details) by controlling for participants’ lack of experience in computer and mobile games. This study differed from most research on overconfidence as we measured performance on multiple occasions to minimise the effect of measurement error. Wayfinding performance recorded at six baseline levels (no. 3, 6, 7, 8, 11 and 12) was used to estimate the magnitude of overconfidence (or underconfidence) for each player in the sample. We then tracked the wayfinding performance of overconfident players at sixteen follow-up levels (game levels no. 13, 16, 17, 18, 21, 22, 23, 26, 27, 28, 31, 32, 33, 36, 37, and 38) to investigate whether their poor performance resulted in early dropout and task failure. We decided to use the specific six baseline levels as they were previously analysed in our recent study (Walkowiak et al., 2023) and therefore we applied a similar approach here to avoid any analytical reproducibility issues and to ensure that this study can serve as a more detailed follow-up of the previous research. Selection of baseline and follow-up levels was additionally validated by regressing performance recorded at follow-up levels on performance at baseline levels. We found that the latter was predictive of wayfinding performance measured at follow-up levels (*t_1,295472_* = 285.01, *p* < .001). Note that levels 4, 8, 12 etc. as well as levels 5, 10, 15 etc. were irrelevant to this study as they employed either a path integration task or purely entertaining game engagement activity (taking a picture of a sea monster), respectively. Other wayfinding levels e.g. 41, 42, and up to level 73, were not utilised as their sample sizes quickly decreased, preventing us from carrying out a reliable country-by-country analysis.

### Measuring wayfinding performance

In order to measure the wayfinding performance at both baseline and follow-up game levels, we have implemented two distance-based performance metrics: the *Standardised and Corrected Distance (SCD)* and the *Mean of Standardised and Corrected Distances (MSCD)*. The former was used to compare wayfinding performance between individual game levels, and it was estimated with Euclidean distance between two-dimensional point coordinates in space sampled at *Fs* = 2Hz for each gameplay. The latter was used to quantify the performance differences between users across several game levels by calculating the arithmetic mean of their respective SCD metrics. Similar approaches were used by Coutrot et al. (2018) and, more recently, Walkowiak et al. (2023), however in the present study we were more interested in individual-level performance rather than general navigation abilities across cultural clusters or countries.

Application of the SCD metric across all six baseline levels to identify overconfident participants allowed us to control for the *“bad luck”* (*i.e.* the measurement error) which is often incorrectly disregarded in studies on overconfidence (2017). Following a recommendation by Coutrot et al. (2018), both metrics were corrected to control for a varying experience in mobile and computer gaming by dividing them by a sum of mean Euclidean distances travelled at levels 1 and 2 in up to three first attempts at each of those two early screening levels. Additionally, the distance metrics were *z*-score standardised to enable unbiased comparison independent of distinct spatial layouts at particular game levels. Finally, both metrics were multiplied by -1 (*i.e.* reversed) to ease the interpretation - the larger the value, the better the wayfinding performance.

In line with the classical approach in research measuring overconfidence, the wayfinding performance in this study was represented as quartiles based on the reversed MSCD metric recorded for the baseline levels and the reversed SCD metric for each follow-up level *i.e.*, players whose reversed MSCD/SCD metric was largest on the particular levels belonged to the 4^th^ quartile (best performance), whereas those with 25% of the lowest reversed MSCD/SCD metrics constituted the 1^st^ quartile (worst performance).

### Measuring overconfidence

The self-reported estimates of participants’ navigation skills were collected during the game after the completion of level 2 via a single question *“How good are you at navigating?”* which offered game players four possible answers: *‘very bad’*, *‘bad’*, *‘good’*, and *‘very good’*. Participants did not have to respond to this question in order to continue playing the game, however our final sample included the gameplays of only these players who answered this question.

To identify overconfident participants, we have subtracted the performance scores estimated at baseline levels (measured with the reversed MSCD metric to avoid the measurement error, and finally converted to the quartiles) from the numeric representation of their self-reported navigation skills (on scale from 1 to 4, where 1 = *‘very bad’* and 4 = *‘very good’*). The resulting difference (*i.e.*, a discrepancy score on numeric scale from -3 to 3) between self-estimated navigation abilities and measured wayfinding performance quartiles was then converted to a categorical variable, where values -3 and -2 indicated *‘underconfidence’*, values between -1 and 1 were considered as *‘normal’* (or accurate estimation of one’s abilities), and values 2 and 3 represented *‘overconfidence’*.

### Definition of cultural clusters

Due to the global scope of data used in this study, one of the objectives was to investigate country-based and cultural variations in wayfinding overconfidence and the magnitudes of the Dunning-Kruger effect. The cultural analysis is particularly problematic as there are several competing methods of establishing clusters based on cultural similarities. However, following Walkowiak et al. (2023), we have adopted the cultural clustering solution implemented in Ronen & Shenkar (2013). It provides the most comprehensive cultural clustering approach which covers a large majority of countries represented in our data and its derivation is based on other influential studies investigating cultural similarities and differences (Hofstede, 2001; Inglehart & Baker, 2000) and the GLOBE project (House et al., 2004). The list of eleven global cultural clusters with their country members is available in Ronen & Shenkar (2013) and Walkowiak et al. (2023). Representatives of countries which were not included in the Ronen & Shenkar’s solution but featured in our data (e.g. Albania, Croatia, Puerto Rico, Saudi Arabia and Vietnam) were removed from the analysis.

### Statistical analyses, covariates, and multicollinearity

The samples defined in the *Data cleaning procedures and sampling* subsection above were used to answer specific hypotheses related to individual differences of overconfidence and Dunning-Kruger effect as well as its impact on dropout with a selection of statistical methods and data reporting approaches.

Firstly, we presented overconfidence ratios by gender, by country and by cultural cluster to identify basic demographic and geographical patterns of overconfidence. Secondly, we implemented a multivariate logistic regression to explain the probability of displaying overconfidence in wayfinding ability with several socio-demographic, environmental and behavioural predictors. Using the same building blocks as in the logistic regression model, we additionally carried out 46 country-level (one for each country) and 11 cluster-level (one for each cultural cluster) logistic regression analyses to investigate how the effects of predictors and their strengths differed between countries and cultures.

To answer the question on significant correlates of the Dunning-Kruger effect, we again presented the ratios of participants (additionally split by gender) who displayed Dunning-Kruger effect (*i.e.*, poor wayfinding performance while being overconfident) on follow-up levels of the game. We implemented level-by-level iterative Pearson’s chi-squared tests with Yates’ continuity corrections to test for statistical significance of differences between female and male proportions who displayed the Dunning-Kruger effect, and then the Bonferroni-adjusted Wilcoxon rank sum tests to estimate significance of difference in age between those overconfident participants who performed well (*i.e.*, no Dunning-Kruger effect) or showed poor navigation abilities (*i.e.*, Dunning-Kruger effect). We then performed 46 country-based (one for each country) and 16 level-based (one for each wayfinding follow-up game level) multivariate logistic regressions to model the probability of the Dunning-Kruger effect by assessing the stability and robustness of independent variables across all countries in the sample and all follow-up levels used in this analysis.

Finally, to answer the research question related to the impact of the Dunning-Kruger effect on task disengagement and game dropout rates, we firstly implemented the Mantel-Haenszel test with the Kaplan-Meier survival estimate to compute the difference in survival between overconfident and not-overconfident users on wayfinding follow-up levels. Secondly, we performed a Cox Proportional Hazards multivariate regression analysis to model the dropout rate with the baseline overconfidence and remaining socio-demographic, environmental and behavioural covariates as independent variables. Finally, to explain the influence of the Dunning-Kruger effect on the early game termination, we implemented a Cox Proportional Hazard multivariate regression with time dependent covariates to account for level-by-level fluctuations in wayfinding performance and time spent on studying maps of each level’s layout.

The models listed above were designed to consider several socio-demographic, environmental and behavioural covariates we collected from the participants. More specifically, we used age and gender of participants as the main demographic independent variables, but our models also included their education attainment (with four levels: *‘no formal’*, *‘high school’*, *‘college’*, and *‘university’*), type of home environment they grew up in (*i.e.*, whether *‘rural’*, *‘mixed’*, or *‘city’*), their average daily commute time (with three levels: *‘up to 30 mins’*, *‘30 mins to 1 hour’*, and *‘1 hour+’*), and average amount of sleep in hours. We used these variables as there has been a growing evidence of their effects on wayfinding performance reported recently with the use of the same data e.g. Coutrot et al. (2022a) showed that those who grew up in low street network environments displayed better navigation performance at SHQ game levels with regular layouts, whereas those who grew up in rural areas or high-entropy cities excelled at levels with irregular layouts. Similarly, the self-reported amount of sleep has been shown to influence spatial navigation - Coutrot et al. (2022b) found that sleep duration is clustered geographically, and it has a non-linear inverted u-shape relationship with wayfinding performance.

We also recorded the duration of viewing a map of level layouts before starting each of the subsequent wayfinding tasks - this variable was used as one of the time-dependent covariates in the final Cox Proportional Hazard model of dropout.

Independent variables in all multivariate models were tested for multicollinearity with the variance inflation factor (VIF) and generalised variance inflation factor (GVIF) using the *car* package (Fox & Weisberg, 2019) for R programming language. The GVIF was used as some terms (e.g. categorical predictors) had degrees of freedom greater than 1. As both VIF and GVIF were close to value 1 across all predictors and models, the results of the models were not due to high correlations between each predictor and the remaining independent variables.

### Data science tools and statistical software

Accessing the full dataset and sampling procedures were implemented with custom-made code written in Python programming language (version 3.9.12). All analyses reported in this manuscript, including data wrangling operations, transformations, statistical tests, survival analysis models and graphical visualisations, were performed in the R language (version 4.2.2) and RStudio Desktop (version 2022.12.10). We used the *tidyverse* family of R packages (Wickham et al., 2019) for data wrangling, transformations, data aggregations and cross-tabulations. Custom-made implementations of native R language were used for inferential hypothesis tests (some of them were also obtained with the *rstatix* package, Kassambara, 2023) and multivariate logistic regression models. The multicollinearity of covariates was tested with the *car* package (Fox & Weisberg, 2019), whereas the survival curves and Cox models were estimated with the *survival* package (Therneau, 2023). Graphical visualisations e.g. survival curves and percentage-at-risk tables were designed with the help of the *ggplot2* (Wickham, 2016) and *survminer* (Kassambara, Kosinski, & Biecek, 2021) packages. All analyses were performed on a Mac Book Pro (2022) with 16 GB of RAM and Apple M2 processor.

## Results

### Identifying correlates of baseline wayfinding overconfidence

The first goal of this study was to identify correlates of wayfinding overconfidence measured at baseline game levels. Based on the estimates of difference between self-reported navigation skills and wayfinding performance quartiles, 28.83% of all participants were identified as overconfident and only 2.57% as underconfident. Self-estimated navigation abilities for the remaining 68.6% of participants closely aligned with their actual game performance. Of those who declared very good navigating skills, almost 44.3% performed below the median (1^st^ and 2^nd^ quartiles) and 70.7% did worse than anticipated (1^st^, 2^nd^ and 3^rd^ quartiles), meaning that only 29.3% of them performed on a par with their self-estimates (4^th^ quartile). On the other hand, 38.3% of those who rated their navigation abilities as very bad displayed very poor wayfinding performance (1^st^ quartile), and only 35.47% of them performed better than the median (3^rd^ and 4^th^ quartiles). Supporting Information, Table S5 displays cross-tabulated proportions of participants by self-estimated navigation skills and wayfinding performance quartiles.

A greater proportion of females than males displayed overconfidence (33.74% vs. 24.69%), but at the same time a slightly larger proportion of women showed underconfidence (2.88% vs. 2.31%; Supporting Information, Table S6). The Pearson’s chi-squared contingency table test between gender and the confidence categories was statistically significant (χ^2^_2_ = 4046.1, *p* < .001).

The between-countries analysis revealed national and cultural differences in overconfidence. For example, in Germany, Costa Rica and Greece, 40.2%, 37.9% and 37.3% of participants, respectively, overestimated their skills, whereas in Ukraine, Finland and Russian Federation this proportion decreased to around 12.4%, 13.3% and 13.4%, respectively. Certain cultural clusters e.g. Germanic (Austria, Germany and Switzerland) and the Near East (Greece and Turkey) countries displayed much higher overconfidence ratios (39.7% and 36.4%, respectively) than the Far East (India, Indonesia, Iran, Malaysia, Philippines, and Thailand), Nordic (Denmark, Finland, Netherlands, Norway, and Sweden) and Confucian (e.g. China, Hong Kong, Singapore, and Taiwan) nations (22.7%, 21.7% and 18.6%, respectively). Also, the Far East and Confucian countries along with players representing Latin Europe (Belgium, France, Italy, Portugal, and Spain) were more likely to record higher underestimation rates of their wayfinding abilities. The proportions of confidence levels by country and cultural clusters are displayed in Figures S1 and S2 of the Supporting Information.

To identify correlates of the wayfinding overconfidence measured at baseline levels, we have initially performed a multiple logistic regression for all participants and countries in the sample. The binary response variable contained values 0 and 1 only: players who underestimated or accurately reported navigating skills were coded as 0 and those who have been identified as overconfident as value 1. The dependent variable was modelled using four categorical (gender, education levels, typical daily commute time, and the type of home environment they grew up in) and two numeric predictors (age and typical average amount of sleep in hours). Independent variables in the model were tested for multicollinearity with the variance inflation factor (VIF) and generalised variance inflation factor (GVIF) as explained in the *Statistical analyses, covariates, and multicollinearity* subsection of Methods, however no high correlations between predictors were reported. The analysis of deviance table using the likelihood-ratio chi-square for generalised linear models revealed that all model predictors were statistically significant with age being the strongest one (*LR*χ^2^_1_ = 39989.69, *p* < .001). The second strongest predictor was gender (*LR*χ^2^_1_ = 3756.25, *p* < .001), followed by the daily commute time (*LR*χ^2^_2_ = 1000.49, *p* < .001). The overall model was statistically significant (*LR*χ^2^_10_ = 45869.18, *p* < .001) but weak as estimated with McFadden’s pseudo-*R^2^* and Nagelkerke *R^2^* (10.13% and 16.4%, respectively). Table 1 presents standardised β coefficients, odds ratios (with 95% confidence intervals) for all terms of the model along with their respective Wald *z* values. According to the model, higher probability of overconfidence was associated with older age, female gender, increased commute time, greater amount of sleep, worse education, and growing up in the city.

**Table 1.** A summary table of the logistic regression model of overconfidence.

| Predictor variables | Standardized $\beta$ Coefficient | Odds Ratio | 95% Confidence Interval | Wald $z$ value |
| --- | --- | --- | --- | --- |
| (Intercept) | -0.602 | 0.55 | 0.52, 0.58 | -23.70*** |
| Age | 0.764 | 2.15 | 2.13, 2.16 | 190.02*** |
| Gender (reference level: *female*) |  |  |  |  |
| *male* | -0.474 | 0.62 | 0.61, 0.63 | -61.12*** |
| Education (reference level: *no formal*) |  |  |  |  |
| *high school* | -0.149 | 0.86 | 0.82, 0.90 | -6.07*** |
| *college* | -0.290 | 0.75 | 0.71, 0.79 | -11.81*** |
| *university* | -0.303 | 0.74 | 0.70, 0.77 | -12.58*** |
| Home environment (reference level: *rural*) |  |  |  |  |
| *mixed* | -0.129 | 0.88 | 0.86, 0.90 | -12.43*** |
| *city* | 0.131 | 1.14 | 1.11, 1.17 | 11.57*** |
| Travel time (reference level: *up to 30 mins*) |  |  |  |  |
| *30 mins to 1 hour* | 0.108 | 1.11 | 1.09, 1.13 | 11.68*** |
| *1 hour+* | 0.311 | 1.37 | 1.34, 1.39 | 31.53*** |
| Sleep | 0.071 | 1.07 | 1.06, 1.08 | 17.74*** |

\* $p < 0.05$ ; \*\* $p < 0.01$ ; \*\*\* $p < 0.001$ .

The direction and strength of the gender variable in the model was a surprising outcome as navigation studies tend to highlight overconfidence of older males (Taillade et al., 2013, 2016; Walkowiak et al., 2023). To investigate it further, we plotted the mean discrepancy score (*i.e.*, defined in the *Measuring overconfidence* subsection in Methods) between the self-estimated abilities and objectively measured wayfinding performance for different gender and age band combinations (Figure S3 Supporting Information). Based on this analysis, it is clear that older men (*i.e.*, 60-70 years old) overestimated more when compared to all other age-gender groups (including all female subsets). This additional analysis provides further support that older males are more overconfident than other individuals, however this finding is not reflected by the linear logistic regression model used here, because it does not capture the non-linearity between predictors and the dependent variable. As a result, the logistic regression model was influenced by the fact that across all other (*i.e.*, younger than 60-70 years old) age groups, females showed higher overestimation than males in the same age bands.

We have additionally carried out logistic regressions for all forty-six countries and eleven distinct cultural clusters represented in the sample (one for each country and cluster) to investigate how the effects of predictors differed between countries and cultures. All country and culture-based models had the same building blocks as the main logistic model defined previously. The McFadden’s pseudo-*R^2^* metrics of obtained models varied between the countries (from 3.6% and 3.7% in China and United Arab Emirates, respectively, to 14.7% and 15.3% in Austria and Chile, respectively) and across cultures (from 3.7% in Arabic, 4.6% in Far East and 5.6% in the Confucian Asian countries to 11.3% for African cluster, 12.4% in Anglo and 14.2% in Germanic nations) suggesting that a single logistic regression built earlier was not a stable enough model to explain cultural differences and varying effects of predictors on overconfidence between countries. Despite this variability, age was still found to be the single strongest positive and statistically significant (based on Wald *z* values, all *p* < .001) predictor of overconfidence across all forty-six countries and eleven clusters. In general, female gender, the greater amount of sleep and longer (over an hour) daily commute time also increased the probability of overconfidence, however these predictors were found to be statistically significant only in certain countries (gender in 41, the amount of sleep in 17, and commute time over one hour in 13 countries out of 46) and cultural clusters (10, 9, and 6, respectively) but not in others. To illustrate the pattern of results, we show in Figure 1 a comparison example of the exponentiated model coefficients (odds ratios) with their 95% confidence intervals for three example countries (Chile, Denmark, and Taiwan) of similar sample sizes (1571, 1660 and 1481 players, respectively) but from distinct cultural clusters (Latin America, Nordic, and Confucian Asia).

**Figure 1.**
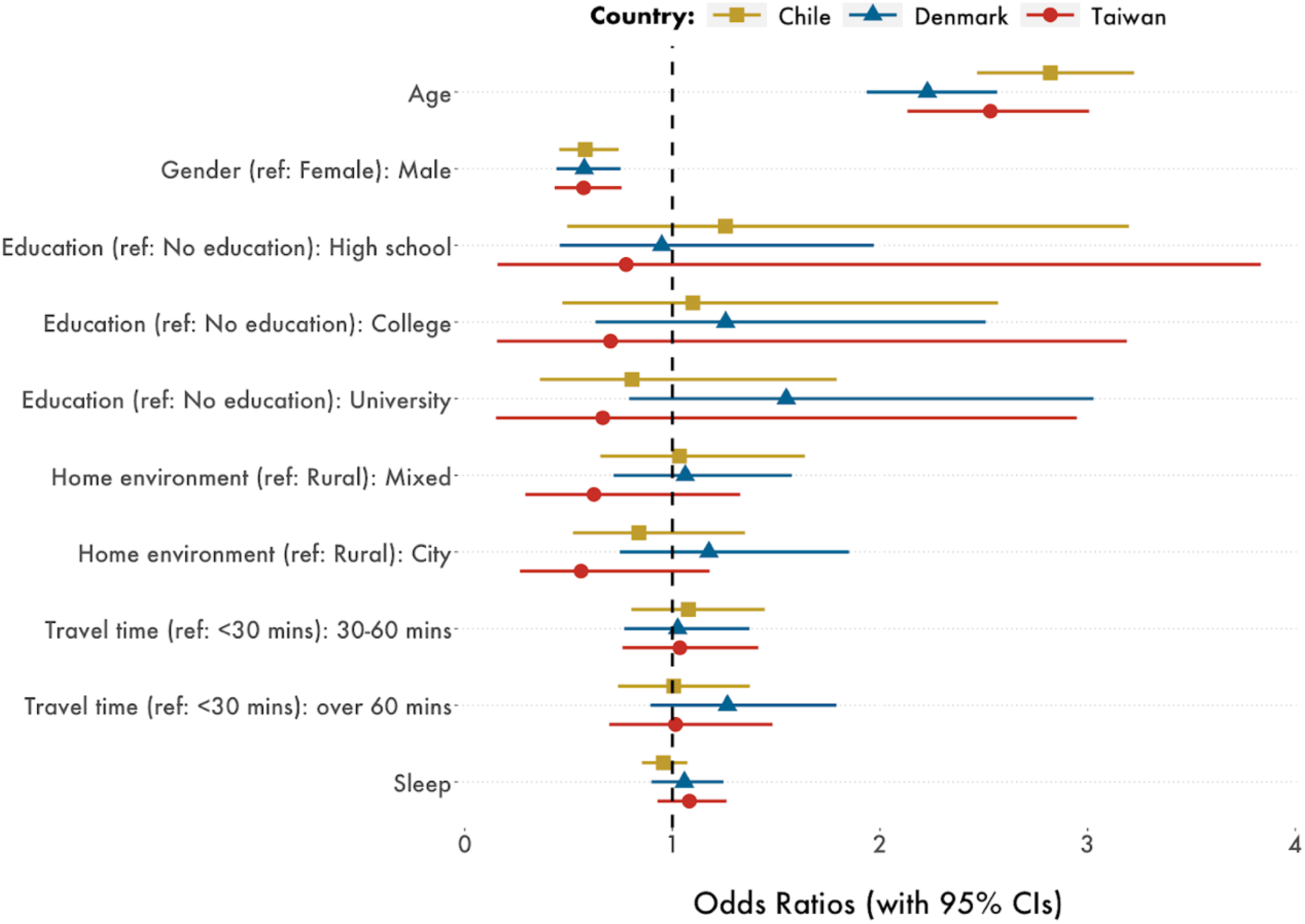
A comparison of country-by-country overconfidence model odds ratios with 95% confidence intervals for Chile, Denmark, and Taiwan. The vertical dotted line separates odds ratios that are greater than 1 (*i.e.*, the overconfidence is more likely to occur as the predictor increases) from those that are smaller than 1 (*i.e.*, the overconfidence is more likely to occur as the predictor decreases). Age was a strong and significant predictor of overestimation in all three countries (and also across all other countries in the sample). The strength and significance of other predictors varied between countries and cultures e.g. the better the education in Chile and Taiwan the lower the probability of overestimation, whereas the increase of education was associated with greater probability of being overconfident in Denmark.

### Who is affected by the Dunning-Kruger bias?

In order to estimate which players were negatively affected by the Dunning-Kruger effect at follow-up game levels, we have extracted a subset of the sample with only those users who completed at least one of the wayfinding follow-up levels and displayed overconfidence at baseline levels (N=86,692). At each wayfinding follow-up level, we have created a new binary variable where players who performed below the median at a specific level (1^st^ and 2^nd^ quartiles) were assigned value 1 (*i.e.*, poor performance, hence Dunning-Kruger effect present), whereas those who performed well and very well (3^rd^ and 4^th^ quartiles) were coded as 0 (no Dunning-Kruger effect). Across all wayfinding follow-up levels, from 62.6% (at level 33) to 67.5% (at level 17) of baseline overconfident players were found to perform below the median at follow-up levels (Supporting Information, Table S7). Greater proportions of females than males exhibited poor performance while being overconfident in all follow-up levels – these differences in proportions were statistically significant as measured with the level-by-level iterative Pearson’s chi-squared tests with Yates’ continuity corrections. Based on the Bonferroni-adjusted Wilcoxon rank sum tests, the overconfident players who performed poorly (*i.e.*, those who displayed the Dunning-Kruger effect) were typically older than those who navigated well (*i.e.*, no Dunning-Kruger effect; Supporting Information, Table S8). The test was not statistically significant only in one out of sixteen follow-up levels (level 33, *p* = 0.28; otherwise, all *p* < .001). The effect sizes for levels where significance was detected were small (*r* between 0.02 at level 31 and 0.17 at level 23).

We carried out sixteen iterative logistic regressions (one for each follow-up level) to model the Dunning-Kruger effect (poor performance while being overconfident; binary dependent variable) with age, map view duration and daily amount of sleep as three numeric predictors and the level of education, gender, home environment, and typical daily commute time as four categorical independent variables. The analysis revealed that older age and female gender significantly increased the probability of the Dunning-Kruger effect across all levels (both predictors with *p* < .001 at each level). Growing up in the city, having worse education, shorter map view duration and longer sleep were identified as significant positive predictors of the Dunning-Kruger effect only at specific game levels (13, 13, 8, and 7 levels out of 16, respectively). The level-by-level logistic regression models were weak, McFadden’s pseudo-*R^2^* metrics varied from 0.7% (for level 31) to 3.5% (for level 32). Finally, we have performed similar iterative logistic regressions for each country. Older age and female gender were again found to be the strongest positive predictors of the Dunning-Kruger effect across all countries – both were significant (*p* < .001) in all sixteen countries. Better education and longer map view duration led to significantly lower probability of the Dunning-Kruger effect in most countries (13 and 11 out of 16, respectively). Longer sleep duration was a positive and statistically significant predictor only in four countries: Czech Republic, Germany, Greece, and Slovakia (*p* = .010, *p* = .021, *p* = .004 and *p* = .003, respectively). Growing up in the city had a statistically significant positive impact on probability of the Dunning-Kruger effect only in two countries (United Kingdom and United States, both *p* < .001), but, on the other hand, in Poland and Greece it was associated with lower probability of the Dunning-Kruger effect (*p* < .001 and *p* = 0.043, respectively). There was no clear pattern of how daily average commute time affected the Dunning-Kruger effect’s probability in both level-by-level and between countries analyses. Similar to the level-by-level data, the country-based models were weak, achieving McFadden’s pseudo-*R^2^* metrics between 1.5% (for Poland) and 3% (in Spain).

### Does the baseline overconfidence or the Dunning-Kruger effect lead to task disengagement and higher dropout rate?

Firstly, we investigated whether the overconfidence estimated at baseline levels was related to the higher dropout rate at wayfinding test levels. Using a sample of 295,474 participants who played all baseline levels and completed at least one wayfinding follow-up level, we have applied the Mantel-Haenszel test with the Kaplan-Meier survival estimate to compute the difference in survival between overconfident and not overconfident users. Table S9 in the Supporting Information presents the level-by-level game dropout ratios for not overconfident and overconfident participants across wayfinding follow-up levels. The test calculations and survival curves were estimated with the *survival* package in R language (Kassambara, 2023). There was a statistically significant difference in survival probability between both groups (χ^2^_1_ = 240.28, *p* < .001). The obtained survival curves (Figure 2) suggested that players who were not identified as overconfident were, in general, more likely to drop out earlier in the game, e.g. only 45.1% of not-overconfident users progressed to level 22 compared to 46.6% of the overconfident group. The progression to level 32 was achieved by 23.8% of the overconfident players, whereas only 21.2% of the not-overconfident users reached the same game level.

**Figure 2.**
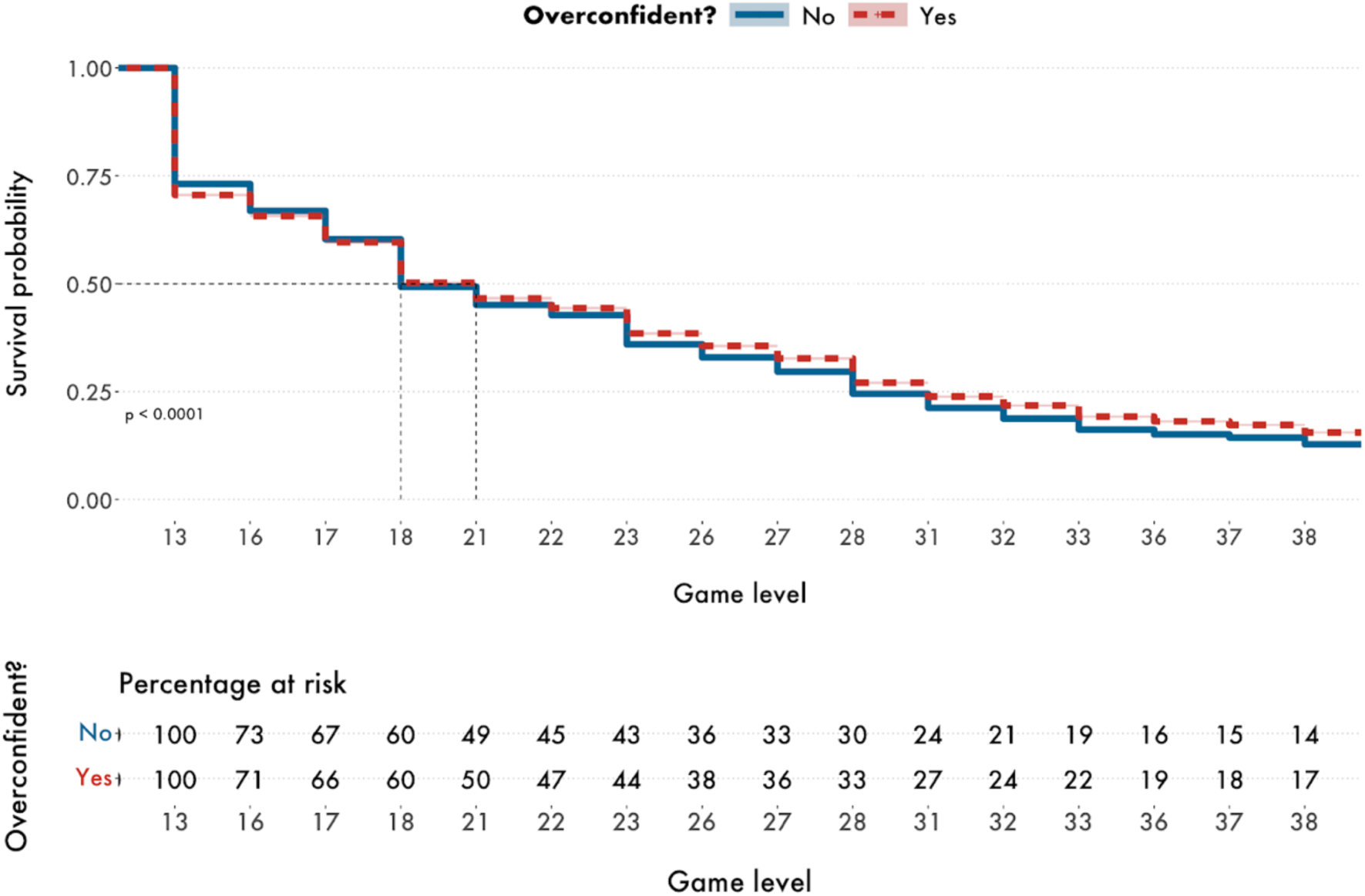
Kaplan-Meier’s survival curves of task dropout for overconfident and not overconfident participants along with the percentage-at-risk table.

Secondly, we have performed a Cox Proportional Hazards multivariate regression model with baseline overconfidence (binary variable), two numeric variables (age and typical amount of sleep), and four categorical variables (education level, gender, daily commute time, and home environment) as predictors. Independent variables in the model were tested for multicollinearity as explained in the *Statistical analyses, covariates, and multicollinearity* subsection of Methods, but no unusually high correlations between predictors were reported. The overall Cox regression model was statistically significant (Wald *z* = 2683, *LR*χ^2^_11_ = 2735, *p* < .001). Based on Wald’s *z* scores, the strongest predictor of task dropout was the daily sleep duration (Wald *z* = -28.51, *p* < .001) - the shorter the average amount of sleep, the higher the risk of dropping out. Being younger, male, having formal education at secondary and tertiary level (*i.e.* high school, college or university), growing up in a mixed environment (*i.e.*, suburban), and reporting shorter daily average commute time were also associated with higher probability of dropout. The overconfidence was found to be a negative significant predictor of game dropout hazard (Wald *z* = -6.76, *p* < 0.001), suggesting that the mere fact of being overconfident (*i.e.*, without taking one’s wayfinding performance into consideration) does not predict early task termination. All standardised terms of the model along with their hazard ratio (*i.e.*, exponentiated coefficients), 95% confidence intervals and Wald *z* values are presented in Table 2.

**Table 2.** A summary table of the Cox Proportional Hazard multivariate regression model of dropout with the baseline overconfidence as one of the predictors.

| Predictor variables | Standardized $\beta$ Coefficient | Hazard Ratio (exponentiated coefficients) | 95% Confidence Interval | Wald $z$ value |
| --- | --- | --- | --- | --- |
| Age | -0.031 | 0.97 | 0.96, 0.97 | -14.70*** |
| Gender (reference level: *female*) |  |  |  |  |
| *male* | 0.061 | 1.06 | 1.05, 1.07 | 15.03*** |
| Education (reference level: *no formal*) |  |  |  |  |
| *high school* | 0.208 | 1.23 | 1.20, 1.26 | 16.04*** |
| *college* | 0.262 | 1.30 | 1.27, 1.33 | 20.24*** |
| *university* | 0.194 | 1.21 | 1.18, 1.25 | 15.19*** |
| Home environment (reference level: *rural*) |  |  |  |  |
| *mixed* | 0.076 | 1.08 | 1.07, 1.09 | 14.51*** |
| *city* | -0.006 | 0.99 | 0.98, 1.01 | -1.04 |
| Travel time (reference level: *up to 30 mins*) |  |  |  |  |
| *30 mins to 1 hour* | -0.050 | 0.95 | 0.94, 0.96 | -10.70*** |
| *1 hour+* | -0.090 | 0.91 | 0.91, 0.92 | -17.41*** |
| Sleep | -0.058 | 0.94 | 0.94, 0.95 | -28.51*** |
| Overconfident (reference level: *no*) |  |  |  |  |
| *yes* | -0.031 | 0.97 | 0.96, 0.98 | -6.76*** |

\* $p < 0.05$ ; \*\* $p < 0.01$ ; \*\*\* $p < 0.001$ .

Both models of baseline overconfidence did not, however, include any information about the level-specific Dunning-Kruger bias. To measure its effect on the dropout rate across the subsequent wayfinding test levels of the game, we have performed a Cox Proportional Hazard multivariate regression with time dependent covariates on the subset of participants who were found overconfident on baseline levels and completed at least one of the wayfinding follow-up levels (N=86,692). A similar approach was used to predict academic dropout by Ameri et al. (2016) and, more recently, to model population evacuation behaviour during hurricanes (Anyidoho et al., 2022). Just like in our models reported previously, two numeric variables of age and typical amount of sleep as well as four categorical variables of gender, education level, average daily commute time, and type of home environment were used as time-independent predictors. Additionally, instead of the binary overconfidence independent variable, we have used two time-dependent level-specific covariates: a.) a previously created binary variable which indicated those who performed poorly (*i.e.*, participants who were affected by the Dunning-Kruger effect; coded as 1) or performed well (coded as 0) at each game level, and b.) the amount of time an individual spent viewing a layout map before each level. No significant and unusually high correlations between predictors were detected. The time-dependent Cox model was statistically significant (*LR*χ^2^_12_ = 843.5, *p* < .001). Poor performance while being overconfident (*i.e.*, Dunning-Kruger effect) was the strongest positive and significant predictor of increased probability of dropping out (Wald *z* =13.67, *p* < .001), whereas the greater amount of sleep, longer time spent on viewing the layout map, longer commute time, having a secondary or tertiary education, as well as being a female statistically decreased the risk of early game termination. It is worth highlighting that age was not found to be a significant predictor of dropping out. All model terms with their standardised coefficients, hazard ratio, their 95% confidence intervals and Wald *z* values are presented in Table 3.

**Table 3.** A summary table of the Cox Proportional Hazard multivariate regression model of dropout with two time-dependent covariates: a level-specific Dunning-Kruger effect and map view duration. The presence of the Dunning-Kruger effect was found to be the strongest predictor of wayfinding task dropout.

| Predictor variables | Standardized $\beta$ Coefficient | Hazard Ratio (exponentiated coefficients) | 95% Confidence Interval | Wald $z$ value |
| --- | --- | --- | --- | --- |
| Age | -0.006 | 0.99 | 0.99, 1.00 | -1.67 |
| Gender (reference level: *female*) |  |  |  |  |
| *male* | 0.022 | 1.02 | 1.01, 1.04 | 3.24** |
| Education (reference level: *no formal*) |  |  |  |  |
| *high school* | 0.142 | 1.15 | 1.11, 1.20 | 7.06*** |
| *college* | 0.180 | 1.20 | 1.15, 1.25 | 8.88*** |
| *university* | 0.137 | 1.15 | 1.10, 1.19 | 6.87*** |
| Home environment (reference level: *rural*) |  |  |  |  |
| *mixed* | 0.070 | 1.07 | 1.05, 1.09 | 7.76*** |
| *city* | -0.010 | 0.99 | 0.97, 1.01 | -1.06 |
| Travel time (reference level: *up to 30 mins*) |  |  |  |  |
| *30 mins to 1 hour* | -0.047 | 0.95 | 0.94, 0.97 | -5.73*** |
| *1 hour+* | -0.089 | 0.92 | 0.90, 0.93 | -10.21*** |
| Sleep | -0.043 | 0.96 | 0.95, 0.96 | -12.45*** |
| Map view duration | -0.057 | 0.94 | 0.94, 0.95 | -10.96*** |
| Dunning-Kruger effect (reference level: *no*) |  |  |  |  |
| *yes* | 0.101 | 1.11 | 1.09, 1.12 | 13.67*** |

\* $p < 0.05$ ; \*\* $p < 0.01$ ; \*\*\* $p < 0.001$ .

Figure 3 presents survival curves from the time-dependent Cox model for four theoretical (*“dummy”*) participants: two females and two males with and without Dunning-Kruger bias, who share the same values for all remaining predictors: arithmetic means of age and map view duration, college education, eight hours of sleep time, daily commuting time of at least one hour and raised in the city (home environment variable).

**Figure 3.**
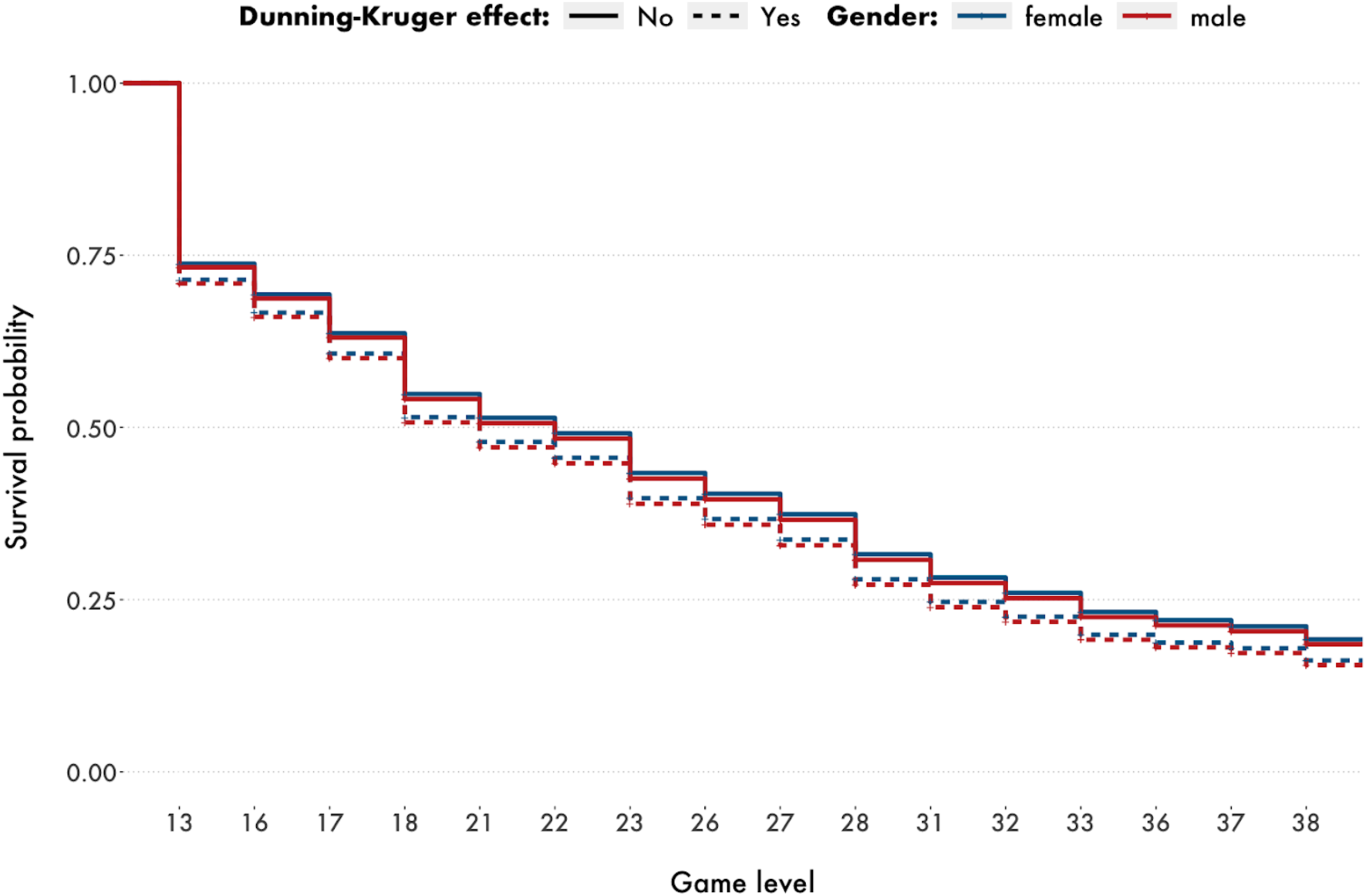
Kaplan-Meier’s survival curves of the dropout risk for four theoretical (“dummy”) participants with varying values of gender and Dunning-Kruger effect variables. Males who displayed high overconfidence and performed poorly were at the highest risk of quitting the game early.

Finally, we investigated whether the Dunning-Kruger effect was predictive of game dropout across all sixteen countries represented in the sample of overconfident participants who completed at least one wayfinding follow-up level. We have implemented sixteen separate (one for each country) Cox Proportional Hazard regression models with the same user demographic predictors and time-dependent, level-specific covariates as in the overall model. The Dunning-Kruger effect was found statistically significant in eleven out of sixteen countries. Slovakia, France, Poland, Brazil and Italy were the only countries where the risk of dropping out was not significantly increased by poor performance amongst the overconfident players, however the direction of effect across all countries was positive - the presence of Dunning-Kruger effect was associated with higher probability of terminating the game early. The prevalence and strength of the Dunning-Kruger bias as a predictor of task dropout may suggest that it was largely independent of geographical location and its impact was relatively stable across different countries. Other important predictors of early game termination in each country generally followed the patterns identified in the overall Cox model applied on the entire sample. Older age, longer sleep time and map view duration were associated with lower risk of dropping out (the first two were found statistically or marginally significant in 11 and 12 out of sixteen countries, respectively, and the map view duration in 10). Also, being a male significantly increased the risk in eight countries, however, in two countries (Canada and United States), male gender significantly lowered the probability of dropout.

## Discussion

This study was the first study focused solely on overconfidence and Dunning-Kruger effect in the field of human navigation and wayfinding measured with a large sample size across multiple countries. Similar to findings from other domains, we reported that, in general, people tend to be more overconfident than underconfident, however, overconfidence may not be as common as some authors claimed in the past. This supports recent evidence which suggests that overestimation is less prevalent across the population and the magnitude of the Dunning-Kruger effect may be smaller than recorded in earlier studies which often did not account for the measurement error or used statistical methods prone to inflating the effect size (Feld, Sauermann, & de Grip, 2017; Gignac & Zajenkowski, 2020). We found that almost 29% of our overall sample overestimated their navigating skills, however it is highly plausible that this rate is dependent on a particular type and specific skill in question. After all, wayfinding, just like walking or reading, is a general skill known and practised daily and effortlessly by most people, whereas more specialised skills e.g. knitting, driving a motorbike or operating machinery are likely to elicit different levels of skill confidence amongst different groups of individuals. Furthermore, we found that the proportions of overconfident people varied between the countries and cultural clusters - Germanic, Near East, African and Latin American countries reported substantially higher rates of wayfinding overestimation than in the Far East, Nordic and Confucian clusters (for further analysis of cross-cultural effects see Walkowiak et al., 2023). This provides additional evidence that the explanation of discrepancy between self-estimates and the objectively measured performance may be more complex than previously thought and requires further research by considering cultural and between-countries variations.

One of the goals of the study was to identify specific predictors of wayfinding overconfidence. We found that older participants and females showed greater tendency to overestimate their skills. The direction of age-related effects aligns with other studies on wayfinding abilities and individual differences (Taillade et al., 2013, 2016; Ulrich, Grill, & Flanagin, 2019, Walkowiak et al., 2023). Although females displayed greater overestimation across most of the life course (19-59 years old), older men (60-70 years old) were prone to much higher overconfidence than all other age-gender groups. This provides further evidence to findings we have recently reported (Walkowiak et al., 2023) that the oldest men are not only more overconfident than younger males, but in fact they tend to overestimate their navigation abilities more than the rest of the population. Other significant factors of navigating overconfidence were longer commuting time, greater amount of sleep, lower level of education and growing up in the city. Although our research was not designed to explain participants’ motivation for their self-reported estimates of navigating skills, it may be that extended travel time to and from work and the greater topographical complexity of areas in which they grew up gave some individuals a false sense of wayfinding expertise which, combined with their lower level of education and higher alertness as a result of longer amount of sleep, might have caused excessively optimistic views of their navigating abilities. Just like the overall rate of overconfidence in the population, the general model of wayfinding overconfidence may be dependent on geographical and cultural factors. We found that certain predictors tend to elicit overestimation to a greater extent in some countries and cultural clusters but not the others. Despite these differences in magnitudes of some factors, we reported that age was the only significant and positive predictor of overconfidence across all countries and cultural clusters.

We were especially interested in investigating the effects of overconfidence coupled with poor performance on the probability of task failure measured by estimating the game dropout rate on multiple follow-up levels. Firstly, we identified those who were more likely to display overconfidence using a set of baseline game levels. We found that significantly greater proportions of females than males exhibited poor performance while being overconfident throughout the game and a set of iterative logistic regressions confirmed that older age and female gender were the strongest predictors of the Dunning-Kruger effect across all game levels as well as all countries in the study. Other factors such as shorter map view duration before each game level, worse education and greater amount of sleep were predictive of poor performance to the lesser extent and in specific countries only. Secondly, we used survival analysis methods to estimate how overconfidence on its own as well as overconfidence together with poor performance (Dunning-Kruger effect) measured as a time-dependent covariate influenced the game dropout rate across follow-up levels. Overconfidence on its own was found to be the weakest of the statistically significant predictors of dropout in the overall hazard model.

Its negative relationship with dropout risk suggested that the increase in overestimation protected individuals from quitting the game early, and it was independent of one’s performance on wayfinding follow-up levels. However, as this model did not account for level-by-level performance fluctuations, it therefore ignored the presence of the Dunning-Kruger effect. When poor performance coupled with overconfidence was used as one of the time-dependent covariates, it became the strongest statistically significant positive predictor of dropout hazard in the overall model, and it remained one of the three most important factors predictive of dropout in eleven out of sixteen countries (it was statistically or marginally significant in fifteen countries). This indicates that the negative consequence of the Dunning-Kruger effect (poor performance while being overconfident) may be responsible for participants’ higher tendency to fail or quit a task early. The strength and presence of this effect across multiple countries indicates that it may exist independently of geographical and cultural differences. Other statistically significant predictors of the overall dropout model such as longer time spent viewing a map before each level, older age, greater amount of sleep, and longer commute time were associated with lower risk of dropping out early. These factors however were less stable between countries than the presence of the Dunning-Kruger effect.

## Data availability

Access to the full Sea Hero Quest dataset (including all wayfinding trajectories generated by the players) is available upon registration through a dedicated server at https://shqdata.z6.web.core.windows.net/.

Please contact the Lead Contacts directly (H.J.S. and E.M.) to obtain the sample of the full dataset used in this study.

Lead Contacts: Hugo J. Spiers and Ed Manley.

Additionally, researchers interested in using the Sea Hero Quest mobile game application can invite participants to play the game and generate wayfinding research data for non-commercial purposes via the SHQ game portal at https://seaheroquest.alzheimersresearchuk.org/.

## Code availability

All data pre-processing operations, statistical analyses and figures included in this manuscript were made using the R and Python programming languages. The custom code scripts are available from the Lead Contacts (H.J.S. and E.M.) upon request.

## Ethics declaration

All participants voluntarily downloaded and played the Sea Hero Quest mobile application game. This study was conducted as part of a larger research project which has been approved by the UCL Ethics Research Committee under the project number: CPB/2013/015. The authors declare no competing interests.

## Author Contributions

Conceptualization, S.W., H.J.S., and E.M. Methodology, S.W, H.J.S., and E.M. Investigation, S.W., H.J.S., and E.M. Formal analysis, S.W., H.J.S., and E.M. Resources, H.J.S., and J.M.W. Data curation, S.W., H.J.S., and J.M.W. Writing - Original Draft, S.W., H.J.S, and E.M. Writing - Review & Editing, S.W., A.C., H.J.S., and E.M. with input from all authors. Visualization, S.W., H.J.S, and E.M. Supervision, H.J.S, and E.M.

## Declaration of Interests

The authors declare no competing interests.

### Acknowledgments

This manuscript was written as part of the Ph.D. research conducted by S.W. whose work was funded by the Alan Turing Institute in London. S.W. would like to thank the Alan Turing Institute for academic support and financial contribution during his doctoral research. Furthermore, the authors wish to thank Deutsche Telekom and Alzheimer’s Research UK (ARUK-DT2016-1) for funding the research and analysis based on the Sea Hero Quest mobile game application. Finally, the authors are grateful to Glitchers Limited for the SHQ game production, and to Saatchi & Saatchi London for the project management and creative input.

