## Supplementary Information for "From overconfidence to task dropout in a spatial navigation test: evidence for the Dunning-Kruger effect across 46 countries"

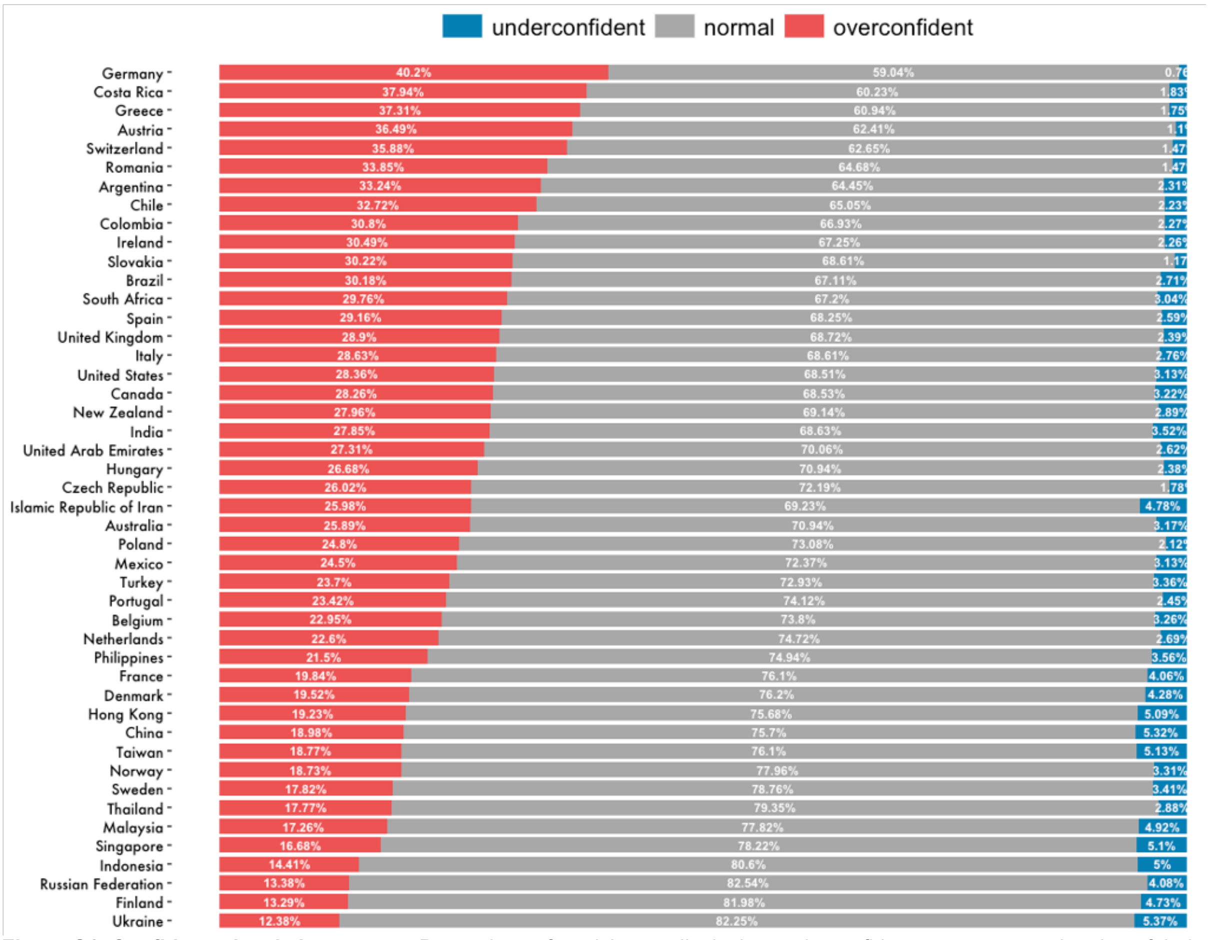


**Figure S1. Confidence levels by country.** Proportions of participants displaying underconfidence, accurate estimation of their skills vs. their actual wayfinding performance, and overconfidence for each of the 46 countries represented in the sample as measured on six baseline wayfinding tasks (levels no. 3, 6, 7, 8, 11, and 12).

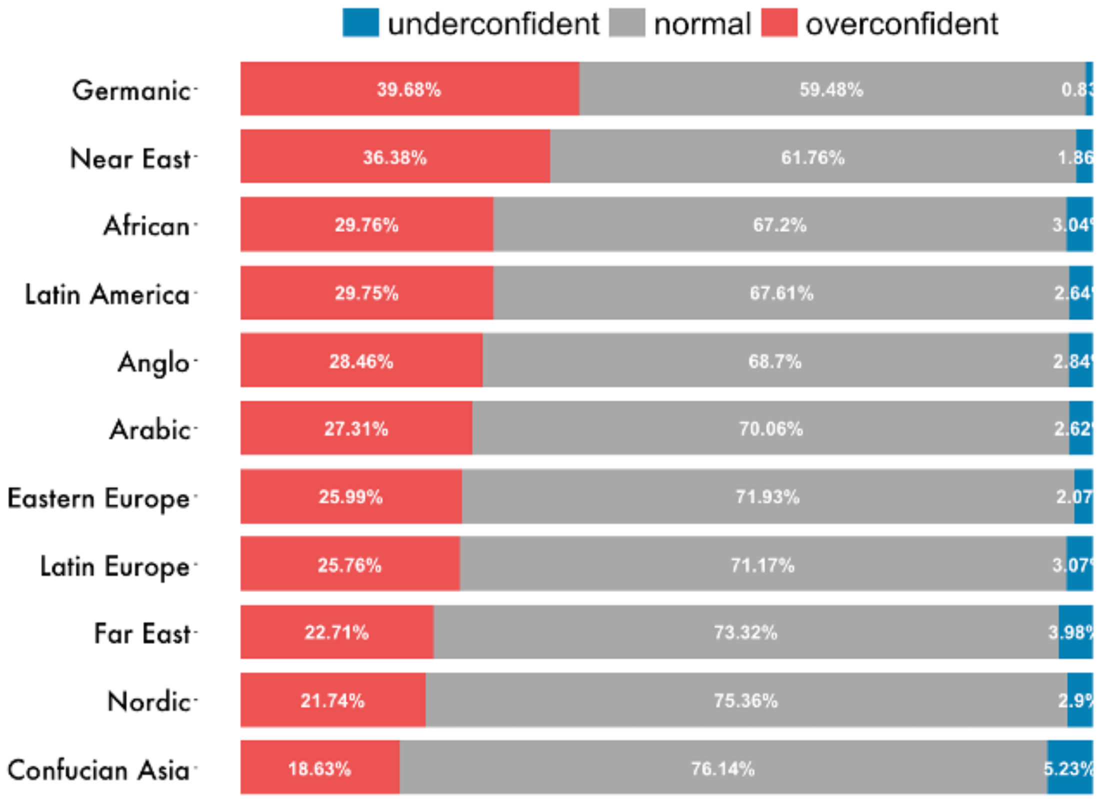


**Figure S2. Confidence levels by cultural clusters.** Proportions of participants displaying underconfidence, accurate estimation of their skills vs. their actual wayfinding performance, and overconfidence for each of the eleven cultural clusters as measured on six baseline wayfinding tasks (levels no. 3, 6, 7, 8, 11, and 12).

**Table S1. Descriptive statistics by country and gender for the sample of 376,836 participants representing 46 countries who played in two practice (SHQ levels no. 1 and 2) and six baseline wayfinding levels (SHQ levels no. 3, 6, 7, 8, 11, and 12).**

| | | | | | Females | | | Males | | |
|---|---|---|---|---|---|---|---|---|---|---|
| | Country | N | Mean age | SD age | N | Mean age | SD age | N | Mean age | SD age |
| 1 | Argentina | 1733 | 34.47 | 12.86 | 671 | 34.98 | 13.81 | 1062 | 34.14 | 12.22 |
| 2 | Australia | 7824 | 39.87 | 14.47 | 4005 | 40.15 | 15.13 | 3819 | 39.59 | 13.73 |
| 3 | Austria | 1902 | 37.20 | 13.49 | 777 | 35.94 | 14.18 | 1125 | 38.06 | 12.93 |
| 4 | Belgium | 1935 | 39.28 | 14.71 | 784 | 40.20 | 15.22 | 1151 | 38.66 | 14.32 |
| 5 | Brazil | 6056 | 33.45 | 12.85 | 2697 | 33.63 | 13.96 | 3359 | 33.30 | 11.88 |
| 6 | Canada | 10666 | 40.72 | 14.87 | 5146 | 40.70 | 15.58 | 5520 | 40.73 | 14.18 |
| 7 | Chile | 1571 | 34.33 | 13.17 | 633 | 33.04 | 13.85 | 938 | 35.20 | 12.63 |
| 8 | China | 4983 | 27.78 | 7.58 | 2513 | 27.05 | 7.33 | 2470 | 28.52 | 7.75 |
| 9 | Colombia | 1409 | 31.68 | 11.41 | 540 | 31.67 | 12.15 | 869 | 31.69 | 10.94 |
| 10 | Costa Rica | 601 | 34.97 | 12.60 | 263 | 35.35 | 13.93 | 338 | 34.67 | 11.46 |
| 11 | Czech Republic | 16201 | 30.36 | 10.84 | 7632 | 30.02 | 11.39 | 8569 | 30.66 | 10.30 |
| 12 | Denmark | 1660 | 37.52 | 13.20 | 558 | 36.60 | 14.07 | 1102 | 37.98 | 12.72 |
| 13 | Finland | 677 | 35.86 | 12.18 | 261 | 34.93 | 12.77 | 416 | 36.44 | 11.77 |
| 14 | France | 7852 | 37.23 | 14.14 | 3301 | 37.60 | 15.21 | 4551 | 36.96 | 13.31 |
| 15 | Germany | 33851 | 41.11 | 13.54 | 15537 | 40.80 | 13.91 | 18314 | 41.37 | 13.21 |
| 16 | Greece | 23773 | 33.34 | 10.74 | 9675 | 32.50 | 10.97 | 14098 | 33.92 | 10.55 |
| 17 | Hong Kong | 962 | 34.46 | 12.11 | 374 | 32.05 | 11.79 | 588 | 35.99 | 12.07 |
| 18 | Hungary | 10422 | 31.70 | 11.42 | 4676 | 31.24 | 11.75 | 5746 | 32.07 | 11.14 |
| 19 | India | 3293 | 26.94 | 8.35 | 507 | 29.47 | 10.92 | 2786 | 26.49 | 7.71 |
| 20 | Indonesia | 1041 | 27.26 | 8.45 | 319 | 26.24 | 7.54 | 722 | 27.71 | 8.79 |
| 21 | Ireland | 1371 | 39.86 | 13.18 | 544 | 39.86 | 14.53 | 827 | 39.87 | 12.23 |
| 22 | Islamic Republic of Iran | 1066 | 32.09 | 9.00 | 286 | 31.21 | 9.66 | 780 | 32.41 | 8.73 |
| 23 | Italy | 11715 | 35.92 | 13.74 | 4637 | 35.08 | 14.06 | 7078 | 36.47 | 13.50 |
| 24 | Malaysia | 1118 | 29.26 | 10.57 | 437 | 28.54 | 10.58 | 681 | 29.72 | 10.54 |
| 25 | Mexico | 3771 | 29.98 | 11.05 | 1439 | 29.39 | 11.31 | 2332 | 30.35 | 10.88 |
| 26 | Netherlands | 22583 | 36.57 | 13.60 | 12386 | 36.71 | 13.96 | 10197 | 36.39 | 13.15 |
| 27 | New Zealand | 1695 | 40.62 | 14.88 | 942 | 40.82 | 15.56 | 753 | 40.37 | 14.00 |
| 28 | Norway | 1089 | 35.46 | 12.34 | 462 | 33.10 | 12.16 | 627 | 37.19 | 12.19 |
| 29 | Philippines | 1321 | 28.24 | 10.14 | 652 | 27.46 | 9.86 | 669 | 29.00 | 10.35 |
| 30 | Poland | 8963 | 29.49 | 9.75 | 4416 | 28.81 | 10.00 | 4547 | 30.16 | 9.45 |
| 31 | Portugal | 2079 | 34.91 | 11.13 | 856 | 33.22 | 11.19 | 1223 | 36.09 | 10.94 |
| 32 | Romania | 3131 | 30.04 | 9.11 | 1125 | 28.50 | 9.09 | 2006 | 30.90 | 9.01 |
| 33 | Russian Federation | 2623 | 27.63 | 7.98 | 905 | 26.35 | 8.50 | 1718 | 28.31 | 7.61 |
| 34 | Singapore | 1295 | 32.77 | 11.63 | 509 | 31.44 | 11.26 | 786 | 33.63 | 11.79 |
| 35 | Slovakia | 4705 | 29.59 | 10.19 | 2110 | 29.20 | 10.47 | 2595 | 29.91 | 9.95 |
| 36 | South Africa | 1119 | 38.58 | 13.70 | 457 | 39.52 | 14.34 | 662 | 37.94 | 13.21 |
| 37 | Spain | 6791 | 37.58 | 13.19 | 2711 | 37.28 | 13.70 | 4080 | 37.77 | 12.85 |
| 38 | Sweden | 1728 | 36.21 | 12.72 | 645 | 34.89 | 12.87 | 1083 | 37.01 | 12.56 |
| 39 | Switzerland | 3052 | 42.49 | 14.15 | 1293 | 41.54 | 14.80 | 1759 | 43.19 | 13.62 |
| 40 | Taiwan | 1481 | 32.46 | 10.74 | 638 | 32.05 | 11.54 | 843 | 32.77 | 10.09 |
| 41 | Thailand | 833 | 31.18 | 11.90 | 317 | 29.19 | 10.82 | 516 | 32.40 | 12.36 |
| 42 | Turkey | 1755 | 31.49 | 9.84 | 580 | 29.67 | 9.79 | 1175 | 32.39 | 9.75 |
| 43 | Ukraine | 614 | 27.29 | 7.82 | 171 | 26.56 | 7.67 | 443 | 27.58 | 7.86 |
| 44 | United Arab Emirates | 648 | 35.76 | 10.57 | 190 | 34.91 | 11.04 | 458 | 36.11 | 10.36 |
| 45 | United Kingdom | 66402 | 43.33 | 14.15 | 29242 | 43.26 | 14.78 | 37160 | 43.38 | 13.65 |
| 46 | United States | 85476 | 39.67 | 14.73 | 43549 | 40.69 | 15.32 | 41927 | 38.61 | 14.00 |

**Table S2. Descriptive statistics by cultural cluster and gender for the sample of 376,836 participants representing 11 cultural clusters who played in two practice (SHQ levels no. 1 and 2) and six baseline wayfinding levels (SHQ levels no. 3, 6, 7, 8, 11, and 12).**

| | | | | | Females | | | Males | | |
|---|---|---|---|---|---|---|---|---|---|---|
| | Cluster | N | Mean age | SD age | N | Mean age | SD age | N | Mean age | SD age |
| 1 | African | 1119 | 38.58 | 13.70 | 457 | 39.52 | 14.34 | 662 | 37.94 | 13.21 |
| 2 | Anglo | 173434 | 41.16 | 14.60 | 83428 | 41.56 | 15.19 | 90006 | 40.78 | 14.02 |
| 3 | Arabic | 648 | 35.76 | 10.57 | 190 | 34.91 | 11.04 | 458 | 36.11 | 10.36 |
| 4 | Confucian Asia | 8721 | 30.05 | 9.79 | 4034 | 28.86 | 9.42 | 4687 | 31.08 | 9.98 |
| 5 | Eastern Europe | 46659 | 30.20 | 10.47 | 21035 | 29.69 | 10.92 | 25624 | 30.62 | 10.08 |
| 6 | Far East | 8672 | 28.52 | 9.60 | 2518 | 28.54 | 10.15 | 6154 | 28.51 | 9.36 |
| 7 | Germanic | 38805 | 41.02 | 13.62 | 17607 | 40.64 | 14.02 | 21198 | 41.34 | 13.26 |
| 8 | Latin America | 15141 | 32.69 | 12.44 | 6243 | 32.64 | 13.35 | 8898 | 32.72 | 11.76 |
| 9 | Latin Europe | 30372 | 36.77 | 13.67 | 12289 | 36.44 | 14.30 | 18083 | 37.00 | 13.22 |
| 10 | Near East | 25528 | 33.22 | 10.70 | 10255 | 32.34 | 10.93 | 15273 | 33.80 | 10.50 |
| 11 | Nordic | 27737 | 36.54 | 13.45 | 14312 | 36.47 | 13.86 | 13425 | 36.61 | 12.99 |

**Table S3. Descriptive statistics by country and gender for the sample of 295,474 participants representing 16 countries who completed two practice (SHQ levels no. 1 and 2), six baseline wayfinding levels (SHQ levels no. 3, 6, 7, 8, 11, and 12) and at least one of the sixteen wayfinding testing levels (SHQ levels no. 13, 16, 17, 18, 21, 22, 23, 26, 27, 28, 31, 32, 33, 36, 37, or 38).**

| | | | | | Females | | | Males | | |
|---|---|---|---|---|---|---|---|---|---|---|
| | Country | N | Mean age | SD age | N | Mean age | SD age | N | Mean age | SD age |
| 1 | Australia | 7012 | 39.96 | 14.48 | 3555 | 40.30 | 15.12 | 3457 | 39.60 | 13.78 |
| 2 | Brazil | 5371 | 33.60 | 12.91 | 2367 | 33.86 | 14.04 | 3004 | 33.40 | 11.95 |
| 3 | Canada | 9603 | 40.71 | 14.90 | 4590 | 40.76 | 15.65 | 5013 | 40.67 | 14.17 |
| 4 | Czech Republic | 15184 | 30.33 | 10.84 | 7187 | 30.05 | 11.41 | 7997 | 30.59 | 10.30 |
| 5 | France | 7155 | 37.38 | 14.18 | 2993 | 37.85 | 15.22 | 4162 | 37.04 | 13.37 |
| 6 | Germany | 30816 | 41.07 | 13.56 | 14108 | 40.80 | 13.92 | 16708 | 41.30 | 13.24 |
| 7 | Greece | 21962 | 33.32 | 10.74 | 8941 | 32.46 | 10.96 | 13021 | 33.91 | 10.55 |
| 8 | Hungary | 9734 | 31.62 | 11.42 | 4383 | 31.17 | 11.76 | 5351 | 32.00 | 11.13 |
| 9 | Italy | 10664 | 36.07 | 13.78 | 4168 | 35.24 | 14.13 | 6496 | 36.61 | 13.53 |
| 10 | Netherlands | 20306 | 36.55 | 13.62 | 11077 | 36.72 | 13.98 | 9229 | 36.34 | 13.18 |
| 11 | Poland | 8305 | 29.49 | 9.79 | 4097 | 28.86 | 10.07 | 4208 | 30.11 | 9.46 |
| 12 | Romania | 2886 | 30.02 | 9.14 | 1037 | 28.48 | 9.12 | 1849 | 30.89 | 9.03 |
| 13 | Slovakia | 4418 | 29.60 | 10.18 | 1990 | 29.21 | 10.49 | 2428 | 29.92 | 9.91 |
| 14 | Spain | 6084 | 37.59 | 13.21 | 2441 | 37.34 | 13.68 | 3643 | 37.76 | 12.88 |
| 15 | United Kingdom | 60430 | 43.40 | 14.16 | 26467 | 43.34 | 14.77 | 33963 | 43.44 | 13.67 |
| 16 | United States | 75544 | 39.69 | 14.77 | 38041 | 40.79 | 15.36 | 37503 | 38.58 | 14.05 |

**Table S4. Descriptive statistics by country and gender for the sample of 86,692 participants representing 16 countries who, based on their performance on six baseline wayfinding levels (SHQ levels no. 3, 6, 7, 8, 11, and 12), were identified as overconfident and completed at least one of the sixteen wayfinding testing levels (SHQ levels no. 13, 16, 17, 18, 21, 22, 23, 26, 27, 28, 31, 32, 33, 36, 37, or 38).**

| | | | | | Females | | | Males | | |
|---|---|---|---|---|---|---|---|---|---|---|
| | Country | N | Mean age | SD age | N | Mean age | SD age | N | Mean age | SD age |
| 1 | Australia | 1800 | 48.94 | 13.17 | 1030 | 48.63 | 13.40 | 770 | 49.35 | 12.86 |
| 2 | Brazil | 1604 | 40.28 | 14.42 | 879 | 39.87 | 15.19 | 725 | 40.77 | 13.43 |
| 3 | Canada | 2705 | 49.75 | 13.64 | 1498 | 49.16 | 14.00 | 1207 | 50.47 | 13.14 |
| 4 | Czech Republic | 3918 | 34.72 | 12.83 | 2278 | 33.83 | 13.07 | 1640 | 35.95 | 12.38 |
| 5 | France | 1409 | 47.69 | 14.00 | 691 | 47.75 | 14.41 | 718 | 47.64 | 13.60 |
| 6 | Germany | 12294 | 47.66 | 12.23 | 6740 | 46.39 | 12.64 | 5554 | 49.21 | 11.54 |
| 7 | Greece | 8126 | 36.62 | 11.34 | 3822 | 34.47 | 11.41 | 4304 | 38.52 | 10.94 |
| 8 | Hungary | 2575 | 36.00 | 12.77 | 1453 | 34.41 | 12.72 | 1122 | 38.05 | 12.53 |
| 9 | Italy | 3052 | 43.38 | 13.78 | 1383 | 41.58 | 14.44 | 1669 | 44.87 | 13.03 |
| 10 | Netherlands | 4542 | 44.57 | 13.23 | 2712 | 44.03 | 13.32 | 1830 | 45.36 | 13.06 |
| 11 | Poland | 2037 | 32.81 | 11.25 | 1186 | 31.71 | 11.37 | 851 | 34.33 | 10.92 |
| 12 | Romania | 974 | 32.55 | 10.03 | 440 | 30.14 | 10.02 | 534 | 34.54 | 9.61 |
| 13 | Slovakia | 1322 | 33.11 | 11.74 | 728 | 31.89 | 11.97 | 594 | 34.60 | 11.30 |
| 14 | Spain | 1762 | 43.91 | 13.01 | 770 | 42.42 | 13.56 | 992 | 45.06 | 12.45 |
| 15 | United Kingdom | 17361 | 50.80 | 12.46 | 8594 | 49.87 | 12.93 | 8767 | 51.72 | 11.92 |
| 16 | United States | 21211 | 49.15 | 13.45 | 12919 | 48.95 | 13.63 | 8292 | 49.48 | 13.15 |

**Table S5. Self-estimated navigation skills by wayfinding performance quartiles.**

| | | Wayfinding performance quartiles (% of sample) | | | |
|---|---|---|---|---|---|
| | Navigation skills | Q1 | Q2 | Q3 | Q4 |
| 1 | very-bad | 38.28 | 26.25 | 19.47 | 16.00 |
| 2 | bad | 32.93 | 26.04 | 22.37 | 18.66 |
| 3 | good | 25.88 | 25.40 | 24.83 | 23.89 |
| 4 | very-good | 20.35 | 23.95 | 26.40 | 29.30 |

**Table S6. Confidence levels by gender.**

| | | Confidence (% of sample) | | |
|---|---|---|---|---|
| | Gender | Underconfident | Normal/accurate | Overconfident |
| 1 | female | 2.88 | 63.38 | 33.74 |
| 2 | male | 2.31 | 73.01 | 24.69 |

**Table S7. The level-by-level ratios of participants who displayed the Dunning-Kruger effect (i.e. poor performance while being overconfident) across sixteen wayfinding testing levels (SHQ levels no. 13, 16, 17, 18, 21, 22, 23, 26, 27, 28, 31, 32, 33, 36, 37, or 38).**

| | Level ID | N | No. of DK effect | DK % |
|---|---|---|---|---|
| 1 | 13 | 86692 | 58207 | 67.14 |
| 2 | 16 | 60739 | 39939 | 65.76 |
| 3 | 17 | 56501 | 38124 | 67.47 |
| 4 | 18 | 51258 | 33798 | 65.94 |
| 5 | 21 | 43085 | 28020 | 65.03 |
| 6 | 22 | 39942 | 25505 | 63.86 |
| 7 | 23 | 38051 | 25028 | 65.77 |
| 8 | 26 | 32914 | 21615 | 65.67 |
| 9 | 27 | 30395 | 20091 | 66.10 |
| 10 | 28 | 27969 | 18148 | 64.89 |
| 11 | 31 | 23066 | 14525 | 62.97 |
| 12 | 32 | 20301 | 13541 | 66.70 |
| 13 | 33 | 18582 | 11624 | 62.56 |
| 14 | 36 | 16395 | 10292 | 62.78 |
| 15 | 37 | 15450 | 9802 | 63.44 |
| 16 | 38 | 14848 | 9645 | 64.96 |

**Table S8. Descriptive statistics for the sample of 86,692 participants who either did or did not display the Dunning-Kruger effect (i.e. poor performance while being overconfident) across sixteen wayfinding testing levels (SHQ levels no. 13, 16, 17, 18, 21, 22, 23, 26, 27, 28, 31, 32, 33, 36, 37, or 38).**

| | | No Dunning-Kruger effect | | | Dunning-Kruger effect | | |
|---|---|---|---|---|---|---|---|
| | Level ID | N | Mean age | SD age | N | Mean age | SD age |
| 1 | 13 | 28485 | 43.31 | 13.88 | 58207 | 46.60 | 14.01 |
| 2 | 16 | 20800 | 42.11 | 14.22 | 39939 | 46.01 | 13.91 |
| 3 | 17 | 18377 | 42.03 | 14.21 | 38124 | 46.16 | 13.90 |
| 4 | 18 | 17460 | 41.90 | 14.42 | 33798 | 46.43 | 13.77 |
| 5 | 21 | 15065 | 43.65 | 14.09 | 28020 | 45.76 | 14.16 |
| 6 | 22 | 14437 | 43.07 | 14.15 | 25505 | 46.33 | 14.03 |
| 7 | 23 | 13023 | 41.91 | 14.13 | 25028 | 46.87 | 13.89 |
| 8 | 26 | 11299 | 42.53 | 14.10 | 21615 | 46.81 | 14.01 |
| 9 | 27 | 10304 | 43.25 | 14.07 | 20091 | 46.57 | 14.13 |
| 10 | 28 | 9821 | 42.81 | 14.37 | 18148 | 47.07 | 13.88 |
| 11 | 31 | 8541 | 44.86 | 14.07 | 14525 | 45.49 | 14.31 |
| 12 | 32 | 6760 | 44.10 | 13.89 | 13541 | 46.18 | 14.38 |
| 13 | 33 | 6958 | 45.51 | 13.95 | 11624 | 45.75 | 14.45 |
| 14 | 36 | 6103 | 44.92 | 14.29 | 10292 | 46.36 | 14.28 |
| 15 | 37 | 5648 | 43.79 | 14.42 | 9802 | 47.24 | 14.09 |
| 16 | 38 | 5203 | 43.93 | 14.37 | 9645 | 47.20 | 14.16 |

**Table S9. The level-by-level game dropout ratios for not overconfident and overconfident participants across sixteen wayfinding testing levels (SHQ levels no. 13, 16, 17, 18, 21, 22, 23, 26, 27, 28, 31, 32, 33, 36, 37, or 38).**

| | | | | | Not overconfident | | | Overconfident | | |
|---|---|---|---|---|---|---|---|---|---|---|
| | Level ID | N | N dropouts | Dropout % | N | N dropouts | Dropout % | N | N dropouts | Dropout % |
| 1 | 13 | 295474 | 81684 | 27.65 | 208782 | 56156 | 26.90 | 86692 | 25528 | 29.45 |
| 2 | 16 | 212521 | 17238 | 8.11 | 151782 | 13011 | 8.57 | 60739 | 4227 | 6.96 |
| 3 | 17 | 195337 | 19014 | 9.73 | 138836 | 13735 | 9.89 | 56501 | 5279 | 9.34 |
| 4 | 18 | 176390 | 31105 | 17.63 | 125132 | 22977 | 18.36 | 51258 | 8128 | 15.86 |
| 5 | 21 | 145082 | 11890 | 8.20 | 101997 | 8769 | 8.60 | 43085 | 3121 | 7.24 |
| 6 | 22 | 133226 | 6922 | 5.20 | 93284 | 4944 | 5.30 | 39942 | 1978 | 4.95 |
| 7 | 23 | 126524 | 19202 | 15.18 | 88473 | 14118 | 15.96 | 38051 | 5084 | 13.36 |
| 8 | 26 | 107245 | 8870 | 8.27 | 74331 | 6355 | 8.55 | 32914 | 2515 | 7.64 |
| 9 | 27 | 98315 | 9415 | 9.58 | 67920 | 6908 | 10.17 | 30395 | 2507 | 8.25 |
| 10 | 28 | 89214 | 15578 | 17.46 | 61245 | 10692 | 17.46 | 27969 | 4886 | 17.47 |
| 11 | 31 | 73588 | 9564 | 13.00 | 50522 | 6781 | 13.42 | 23066 | 2783 | 12.07 |
| 12 | 32 | 64083 | 6985 | 10.90 | 43782 | 5197 | 11.87 | 20301 | 1788 | 8.81 |
| 13 | 33 | 57270 | 7572 | 13.22 | 38688 | 5350 | 13.83 | 18582 | 2222 | 11.96 |
| 14 | 36 | 49736 | 3326 | 6.69 | 33341 | 2359 | 7.08 | 16395 | 967 | 5.90 |
| 15 | 37 | 46518 | 2209 | 4.75 | 31068 | 1503 | 4.84 | 15450 | 706 | 4.57 |
| 16 | 38 | 44535 | 4772 | 10.72 | 29687 | 3253 | 10.96 | 14848 | 1519 | 10.23 |
